# EoRNA2: Autonomous Data Discovery and Processing for Databasing of Gene Expression Data

**DOI:** 10.64898/2026.03.11.710847

**Authors:** Linda Milne, Craig G. Simpson, Wenbin Guo, Claus-Dieter Mayer, Iain Milne, Micha Bayer

**Affiliations:** Dept of Information and Computational Sciences, The James Hutton Institute, Invergowrie, Dundee, DD2 5DA, Scotland, UK; Dept of Cell and Molecular Sciences, The James Hutton Institute, Invergowrie, Dundee, DD2 5DA, Scotland, UK; Biomathematics and Statistics Scotland, University of Aberdeen, Aberdeen, AB25 2ZD, Scotland, UK; Higentec Breeding Innovation (ZheJiang) Co., Ltd., Lishui, 323000, China

## Abstract

We describe a major new release of the EoRNA database, a gene expression database for barley based on public data, first published in 2021^1^. EoRNA v.2 (https://ics.hutton.ac.uk/eorna2/index.html) features an order of magnitude more samples and is based on a new automated workflow of sample discovery and processing which has enabled a dramatic scale-up the original database. EoRNA v.2 also features a major rebuild of the web user interface with rich new functionality. All infrastructure-related code and database schemas and web components are now species agnostic and publicly available for reuse with other taxa. A dedicated new reference transcript dataset has been created for EoRNA v.2 which is largely based on the recently published barley pan-transcriptome and represents the most comprehensive dataset of its kind to date.

## Background

The growth of publicly available short read sequencing data shows no sign of abating, with the total volume of short read data in the European Nucleotide Archive at 94 petabytes at the time of writing (https://www.ebi.ac.uk/ena/browser/about/statistics). However, the reuse of publicly available datasets is limited^2^ which means a missed opportunity in terms of the scientific value of this vast data resource. This includes the potential for reanalysis of RNA-Seq data which can be used to quantify gene expression. Efforts have been made to quantify large volumes of public RNA-Seq data from human and mouse^3^ ^4^ or other model organisms^5^ but there is still little reuse of this type of data in the plant science community. The EoRNA2 database and website described here for barley provides an automated pipeline for the analysis and presentation of gene and transcript data for any species. Barley is a valuable cereal crop used largely for animal feed, human nutrition and the production of malt for beer and spirits^6,7^. It has a long domestication history, identified as one of the founding crops with evidence of multiple domestication events^8–10^. Barley has a diploid genome (2n = 14) and established reference sequence (∼5.1Gb) as well as pan-genomes, which allows simpler genetic mapping and gene analysis compared to its polyploid relatives^11–16^. There are extensive collections of wild barley, landraces, mutants, near-isogenic lines, TILLING and segregating mapping populations used for quantitative trait linkage analysis, genome wide association studies, nested association mapping and breeding^17–24^. Barley is further amenable to transformation and gene editing to facilitate functional testing of selected genes^25,26^. Barley is a principal model crop for the study of temperate cereals^27^ and has a broad genetic diversity that allows selected cultivars and landraces to produce yield in semi-arid/arid areas and at high altitudes and high longitudes, where seasons can be short and cold^22,28–32^. Consequently, barley has been used for studies in photoperiod and flowering time regulation as a response to growth in different latitudes and environments, and studies of seed development and grain quality^33–35^. Barley has natural tolerances and environmental adaptations that have been utilised to develop grain improvement, stress resistance and environmental adaptations in other cereals either through gene identification, markers or direct gene transfer^28,36–38^. Many gene expression studies have, therefore, been conducted in barley to identify genes in biochemical pathways that respond to the environment and its stresses, and to characterise the changes in barley throughout its growth and development.

## Methods

### Generation of the EORNA2 reference transcript dataset

In order to allow most transcripts present in the public RNA-Seq data to be mapped successfully, the most complete set of reference transcripts possible was prepared for mapping. A number of barley reference transcriptome datasets (RTDs) have been published in recent years, and we chose to combine three RTDs, which we reasoned should capture a large proportion of total barley transcript diversity between them:

1. BaRTv2^39^, a recent single-cultivar RTD based on the cultivar Barke, featured highly accurate transcripts generated using a long-read-based assembly as the backbone, complemented by short-read assemblies to enhance splice junction accuracy and transcript diversity. This RTD was included principally for backward compatibility and transcript quality.
2. Morex RTD (HvMx)^40^ was produced as part of the barley pan-transcriptome project^40^ and featured a rich array of biotic and abiotic stress genes from the barley reference cultivar Morex, which were less well represented in BaRTv2.
3. PanBaRT20^40^, the primary barley pan-transcriptome resource, covered five different tissue types and 20 different genotypes, and thus provided genotypic diversity as well as a comprehensive catalogue of tissue-specific genes, but did not explicitly target stress response genes. This RTD was also assembled using the hybrid approach that integrates both long-read and short-read sequencing data.

Transcript sequences from BaRTv2 and Morex RTD (HvMx) were mapped with Minimap2^41^ to PSVCP20, the linear pan-genome sequence produced as part of the barley pan-transcriptome project^40^ and merged using a custom script (https://github.com/cropgeeks/eorna-v2/blob/main/scripts/RTD/mergeRTDs.r). This linear pan-genome sequence also serves as the reference genome of PanBaRT20, with all tissue and genotype samples from the pan-transcriptome project having been mapped to this and is described in more detail in the original publication^40^. Minimap2 was used in splice mode (-ax splice:hq) and the maximum intron length size was set to 15,000 base pairs (bp) (-G 15000). Secondary and supplementary alignments were removed. Samtools^42^ was used to filter the resulting SAM files, and for conversion to BAM format as well as sorting of BAM files. BEDTools^43^ and UCSC utilities^44^ were used to convert the BAM files to GTF format. The GTF files of BaRTv2 and Morex RTD, mapped to the linear pan-genome, were merged with the GTF file of PanBaRT20. Overlapping transcripts were assigned to the same gene ID. To reduce redundancy, mono-exon and multi-exon transcripts were processed separately. Mono-exon transcripts within the same gene were collapsed into a single representative transcript, with start and end positions defined by the minimum and maximum coordinates of the original mono-exon transcripts. Multi-exon transcripts were grouped by intron combinations, with transcripts sharing identical intron structures considered redundant. These were merged into a single representative transcript, with its start and end positions determined by the minimum and maximum coordinates of the grouped transcripts. Intronic genes entirely contained within the introns of other genes were annotated with distinct gene IDs. Chimeric genes that overlapped with more than one gene locus were identified based on their overlapping regions. When a set of transcripts could be assigned to two gene models, and the overlap between these two models was <5% of each gene model’s length, these models were assigned different gene IDs.

The merged RTD was exported in GTF format for subsequent analyses. Transcript sequences were extracted from the linear pan-genome using gffread^45^, generating a FASTA file representing the final RTD for downstream analyses, such as functional annotation and expression analysis.

The approach described here ensures comprehensive integration of transcript data across different barley reference datasets, reducing redundancy and enhancing annotation quality for the EoRNA2 RTD. All scripts used for this workflow are available at https://github.com/cropgeeks/eorna-v2/blob/main/scripts/RTD.

### Functional annotation of the RTD

The EoRNA2 RTD transcript sequences were functionally annotated with three separate, complementary approaches to maximise the number of transcripts receiving annotation. We first annotated transcripts using the web-based annotation tool TRAPID^46^ (https://bioinformatics.psb.ugent.be/trapid_02/). We extracted the longest transcript for each gene as representative and submitted these for online annotation. Output files were combined with a custom script that merges GO and Interproscan annotation with transcript metadata full length information.

For the two protein-based annotation methods we first translated the transcripts using Transuite^47^ on default settings. We then used Pannzer^48^ (http://ekhidna2.biocenter.helsinki.fi/sanspanz) to generate annotations for the longest protein of each gene. We used a custom script to parse the Pannzer output for downstream merging with other annotation data.

The third annotation method used was AHRD (Automated Assignment of Human Readable Descriptions, https://github.com/groupschoof/AHRD), a text mining tool that generates human readable functional descriptions from the output of one or more BLAST^49^ searches. We used the longest proteins as queries in BLAST searches against the Swissprot and TREMBL databases^50^ and then ran the outputs through the AHRD tool. Finally, we combined the outputs from all three annotation methods by gene ID, using a custom R script. Despite using multiple annotation approaches, 33.8% of genes did not have any functional annotation (Table 1), most of these non-protein coding transcripts that had not been annotated. All scripts used for annotation are available online at https://github.com/cropgeeks/eorna-v2/tree/main/scripts/RTD/annotation.

**Table 1.**
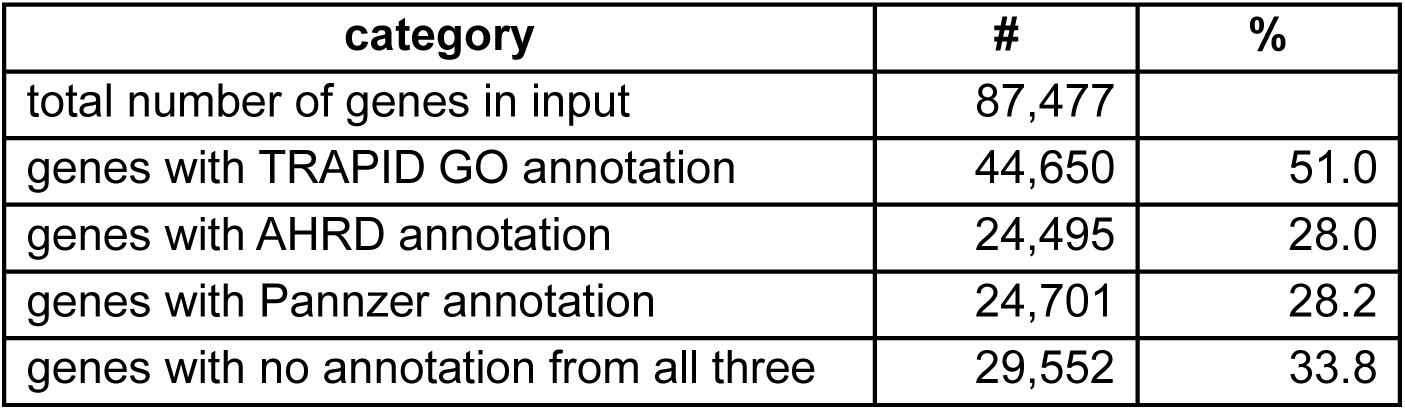
Annotation statistics for the EORNA2 RTD.

| category | # | % |
| --- | --- | --- |
| total number of genes in input | 87,477 |  |
| genes with TRAPID GO annotation | 44,650 | 51.0 |
| genes with AHRD annotation | 24,495 | 28.0 |
| genes with Pannzer annotation | 24,701 | 28.2 |
| genes with no annotation from all three | 29,552 | 33.8 |

### Workflow for data discovery, download and quantification

To facilitate easy, reproducible data processing, we fully automated the process of discovery of relevant RNA-Seq studies and quantification of their sequence read data. We used the REST (Representational State Transfer) API (Application Programming Interface) at the European Nucleotide Archive (ENA) for the purpose of both study and sequencing run discovery (https://ena-docs.readthedocs.io/en/latest/retrieval/programmatic-access.html). The ENA mirrors all short-read data from the other INSDC (International Nucleotide Sequence Database Collaboration)^51^ partners, and so a query to any one of the three repositories will result in the same set of accessions. We implemented a single Nextflow^52^ workflow that generates both study and run queries and then proceeds to download and process the data (Fig 1). The detailed sequence of steps is as follows:

1. Given one or more NCBI taxon identifiers, send a programmatic query to the ENA’s REST API to retrieve a list of all project accessions that contain paired end RNA-Seq for our species of interest.
2. For each of the projects retrieved, send another programmatic query to the ENA’s REST API which retrieves a list of that project’s run accessions and their metadata. Add these to a combined metadata file.
3. For each run accession, download the raw FASTQ files from the ENA, given the URLs in its metadata.
4. For each pair of downloaded FASTQ files, remove poor quality bases and adapter sequences from the reads using the trimming tool fastp^53^. This also produces a QC report.
5. For each pair of trimmed read files, conduct gene expression quantification using the tool Salmon^54^ against the EoRNA2 RTD.

**Figure 1.**
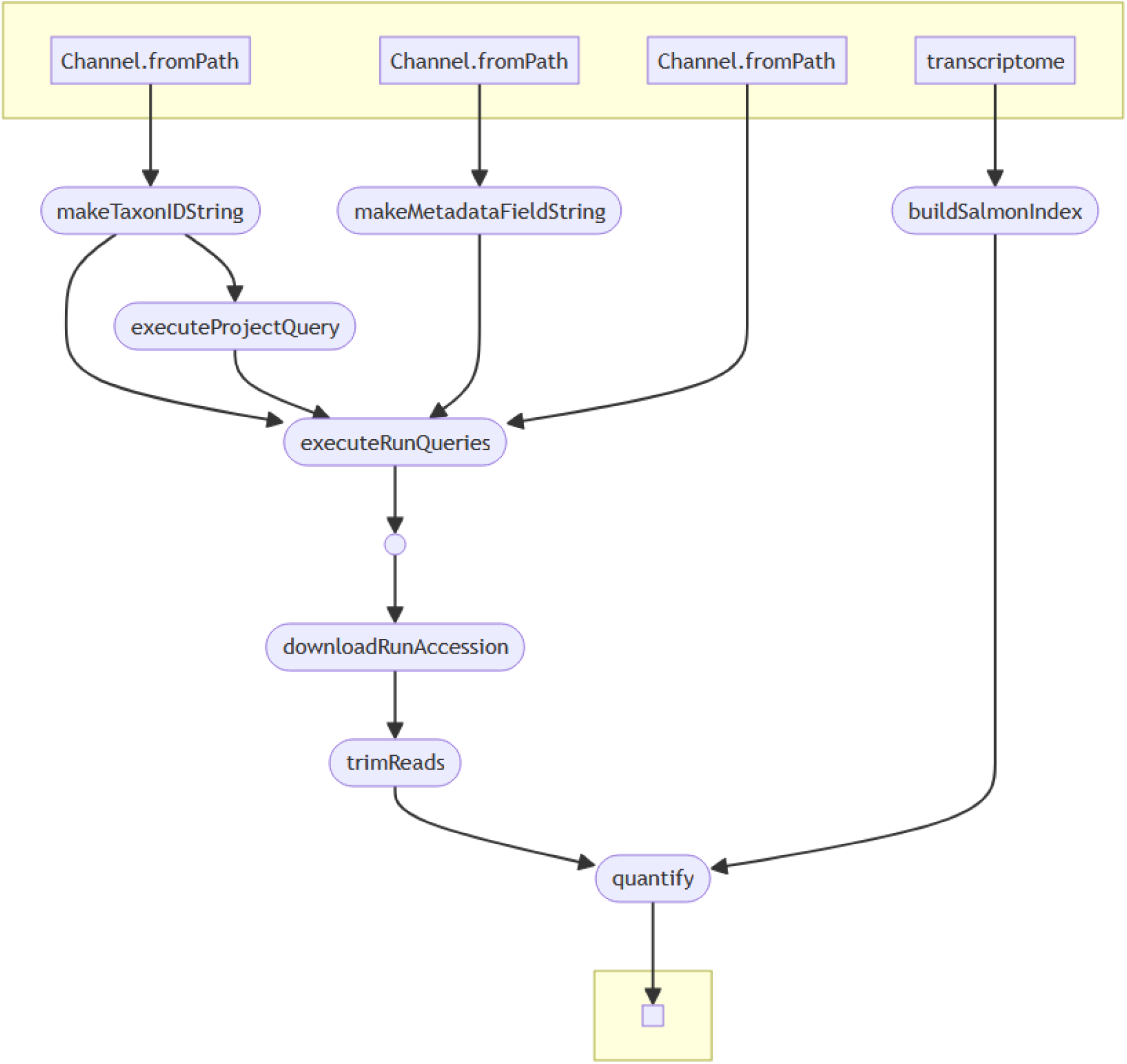
Nextflow workflow implementation for data discovery and processing.

The workflow requires minimal inputs and no software installation (except Nextflow itself) as this is handled by Nextflow’s dependency handling protocol which allows dynamic deployment of software at runtime. For this we opted for the Biocontainers^55^ option which supports most of the commonly used bioinformatics packages. These are downloaded at runtime as Docker^56^ images and require no installation. Inputs required for the workflow include a FASTA file with reference transcript sequences for the quantification step and one or more NCBI taxon identifiers (https://www.ncbi.nlm.nih.gov/guide/taxonomy/) that define the taxa for which RNA-Seq is to be collected. Nextflow supports multiple execution environments, ranging from local execution on a single machine to common High Performance Computing (HPC) scheduling systems such as Slurm^57^ and PBS^58^, to commercial cloud computing services such as Amazon Web Services.

Based on extensive trial runs from our EoRNA v.1 database^1^ we had a comprehensive catalogue of error conditions that can occur at various stages of the workflow. We implemented appropriate error handling strategies for each of the commonly encountered scenarios. Temporary server failures, due to e.g. too many simultaneous requests, are handled through Nextflow’s retry strategy which allows for a defined number of retries, in this case in conjunction with an appropriate wait period. Other errors such as misconfigured metadata (e.g. three instead of two download URLs) are handled within the subtask code and result in specific error codes being emitted for downstream troubleshooting purposes.

The workflow is available for download from https://github.com/cropgeeks/eorna-v2/tree/main/scripts/quantification/nextflow.

### Construction of the EORNA2 database

The database and website were made following the methods used for the original EoRNA v1 database^1^, with the exception of using CanvasJS (https://canvasjs.com/) to draw the interactive TPM plots instead of Plotly (https://plotly.com/r/). This change was introduced to speed up the rendering of the graphs due to the much larger number of samples and larger number of transcripts per gene in this new version of the database. Further changes include some alterations to the database structure and CGI/Perl scripts to improve the speed of data retrieval and page rendering.

The website interface allows users to search for a gene of interest via a nucleotide or protein sequence, by a known gene ID, and by keyword search of the gene annotation. In this new version of EoRNA the pangenome may be explored through the “Region Search” page which offers the user the option to obtain a list of all genes in a region of a chromosome, either by using a specified position or by naming two gene IDs on a chromosome. The returned list of genes allows the user to then see the genes within the integrated JBrowse^59^ genome viewer which then provides links to explore the expression values of the EoRNA2RTD genes.

The new version includes an updated set of eight barley transcriptomes, which have been compared to the EoRNA2RTD so that a user can discover a related EoRNA2 gene using a gene ID from one of the other transcriptome datasets. The correlation between the nine barley transcriptomes was made using GMAP^60^ to find the location of all the transcripts on the linear pan-transcriptome sequence. GffCompare^45^ was then used to find the co-located transcripts. All transcriptome relationship data was stored in the database and gene locations were added to the JBrowse genome viewer.

Due to the large number of samples included in the database, users can select subsets of samples by filtering based on sample metadata, such as cultivar, tissue type, experimental conditions etc. Every study included has had appropriate keywords added to help guide sample choice for inclusion in the TPM graph.

Perl scripts have been provided in our github repository (https://github.com/cropgeeks/eorna-v2) to create SQL files for the MySQL database. Users need to provide the transcriptome both as a GFF file and as a FASTA file, and the Salmon output files. A tab-delimited file of the metadata for each sample is also required. The website can be created using the provided script by adding the user’s preferences for the website name etc, and the name of the MySQL database. The scripts are available on the project github page (https://github.com/cropgeeks/eorna-v2) along with instructions on how to use them.

## Data Overview

### EoRNA2_RTD, an extended barley reference transcript dataset

Extensive barley RNA-Seq datasets are deposited in public repositories that portray barley gene expression across daily circadian growth cycles, different tissues and organs, environmental adaptation and genotypic variation. This has led to the establishment of barley transcriptomes, reference transcript datasets (RTDs) and gene expression atlases visualised through interactive websites that summarise gene expression responses across these cell types and growth conditions ^1,40,61–64^. However, these resources feature a limited subset of the available studies for barley and, except for EoRNA v.1, are based on reference sequences with low transcript-level resolution. RTDs have improved through the combination of short-/long-read sequencing and refined statistics, leading to well defined splice junctions and transcription start and end sites^39,40,65^, whilst at the same time quantification algorithms such as Salmon and Kallisto have continued to evolve and improve, e.g.^66^. Previous barley RTDs have been assembled using a limited number of studies that do not account for the extensive gene and transcript variation that barley genotypes hold, or the alternative transcripts produced by post-transcriptional processes. The first EoRNA transcript abundance database uniquely showed transcript level expression from the BaRTv1.0 RTD represented by 60,444 gene models and 177,240 transcript sequences (https://ics.hutton.ac.uk/eorna/index.html) ^1^. This was followed by firstly BaRTv2, a more accurate barley RTD from cultivar Barke, consisting of 21 tissues, growth stages and treatments^39^, secondly, a database describing the expression of 20 genotypes across 5 different barley tissues (https://ics.hutton.ac.uk/panbart20/index.html) and thirdly, a genotype-specific (cv. Morex) database across 12 different Morex experiments and 315 replicated samples (https://ics.hutton.ac.uk/morexgeneatlas/index.html), both part of the development of a barley pan-transcriptome^40^. We have merged these latter three RTDs to create EoRNA2_RTD, the most complete barley RTD assembled so far, which contains 87,476 genes and their variants and 653,285 transcripts. EoRNA2_RTD was assembled against the linear pan-genome assembly of the 20 genotypes of the first barley pan-genome^67^ and showed an average mapping rate of 89.2% (Fig. 2).

**Figure 2.**
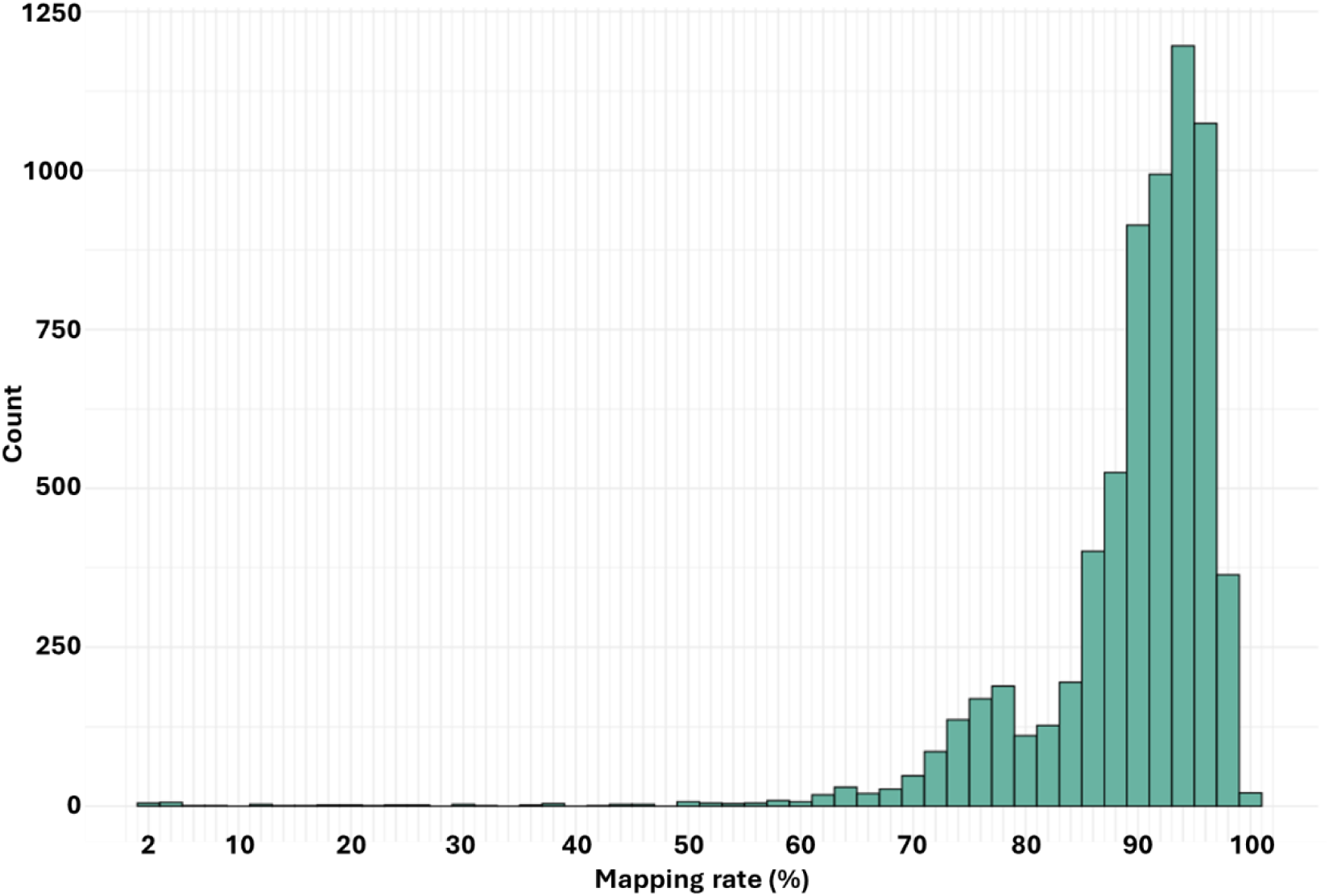
Distribution of mapping rates for all barley RNA-Seq samples analysed in this study. The mapping rate is the percentage of reads in the input data that mapped to a transcript in EoRNA2_RTD.

### EoRNA2 displays transcript variation

EORNA v.2 features a total of 171 studies comprising 6,285 sample accessions. This represents the full complement of paired-end Illumina RNA-Seq from barley in the European Nucleotide Archive as of May 2024. The barley EoRNA2 database uniquely showed transcript variation in expression across its 653,285 transcripts. It was estimated that around 80% of plant genes were multi-exonic and that around half of these genes generated multiple alternative transcripts by alternative pre-mRNA splicing^68,69^. Without a change in transcription rate, alternative splicing may vary with different tissues and conditions, leading to genes that show differential alternative splicing (DAS) of transcripts and in some cases, transcript isoform switching that is the result of a change in abundance of one transcript for another in contrasting samples^70–72^. EoRNA2_RTD also displayed transcript models that showed alternative transcript starts and stops that have been shown to be important in gene regulation^73,74^. Transcript abundances are, therefore, important to determine differential expression responses, but are rarely portrayed in gene expression studies. EoRNA2_RTD included genes and their transcripts that were the result of transcriptional and post-transcriptional processes that allow quantification and presentation of both gene and gene transcript expression. Finally, EoRNA2_RTD displayed genotype sequence variants that may alter the protein coding sequence or lead to changes in splice sites through mutation.

### Gene expression quantification by Salmon

Barley RNA-Seq data continues to expand exponentially from the 22 RNA-Seq studies (843 RNA-Seq samples) available in 2021^1^, to the 171 RNA-Seq studies (6,285 RNA-Seq samples) presented in EoRNA2. Fast and accurate quantification of short-read RNA-Seq data can be achieved with pseudo-alignment bioinformatics tools such as Kallisto and Salmon^54,75^. In the production of EoRNA v.1, the accurate normalisation across all the samples occurred using Salmon^1^, and thus Salmon was also used for quantification of the barley RNA-Seq datasets for EoRNA v.2. The gene expression quantification output from Salmon quantification uses Transcripts Per Million (TPM) as units and gives a normalised measure of gene expression that is comparable both within single and across multiple samples, by accounting for transcript length and sequencing depth in that order. We quantified all 6,285 downloaded short-read sequence samples across the 171 different experiments using EoRNA2_RTD and present the numerical expression outputs in an easily accessible database and website (https://ics.hutton.ac.uk/eorna2/index.html). Numerical quantification values are provided at the transcript level allowing the visual presentation of both gene and transcript expression levels, suitable for both differential gene expression (DGE) and differential alternative splicing (DAS) studies. The EoRNA v.2 database and website represents the most comprehensive gene and transcript expression dataset described so far for barley. It provides archived barley RNA-Seq datasets in a format that the research community can interrogate and use for their research areas and gene discovery. Gene and transcript expression data is linked to sequence information, barley gene synonyms, best match homologies to Arabidopsis, rice and Brachypodium, and is also linked to Gene Ontology (GO) annotation.

The primary value of comparing between sample sets is that it allows researchers to answer how their gene of interest is expressed in specific tissues or under which environmental conditions. Here, individual transcript RNA-Seq expression results are displayed in graphical form, as TPM values directly from the output of Salmon. These interactive plots, therefore, simply permit rapid visual assessment of expression levels in a selected gene of interest. To validate the data, we present examples of genes that illustrate the advantages and limitations of this broad range of barley expression data.

### Metadata curation

The metadata accompanying the read data was subject to extensive manual curation which identified several samples, and in some cases entire projects (n =25), that were unsuitable for processing. We identified recurrent issues with the configuration of the metadata which were as follows:

- No enforced standardisation of metadata required
- Studies include taxa other than barley
- Empty fields in the downloaded metadata table
- Metadata added in an inappropriate table field
- Samples split across more than one run accession
- Samples have one or three FASTQ files listed, not a pair
- Discrepancy between the study accession number in the metadata table and the published paper
- Spelling errors and inconsistency of nomenclature between samples from the same study
- Duplicate studies with different study accession numbers

For basic functionality we expected sample metadata to include the cultivar, tissue type, developmental stage, experimental conditions, and biological replicate number. Often this meant finding the published paper associated with the study and using the methods and materials described to fill in empty fields. Samples for which no clear metadata was available were not included. We encountered substantial diversity in the description of experimental conditions (e.g. cold stress was described in different ways), so appropriate keywords were given to each sample.

## Technical Validation and Use

### TPM variation across photosynthetic and non-photosynthetic samples

One of the aims of the EoRNA2 database is to present differences in expression across multiple experimental samples. While this is highly intuitive when gene expression is specific only to the tissue/organ or condition (see below) it may be less so where genes are expressed across a range of different tissues as presented in EoRNA2. This is particularly the case in plant gene expression studies with comparisons between genes expressed specifically in photosynthetic tissues (e.g. leaf) compared to tissues that do not express these genes, e.g. root (Figure 3A). Using the root and shoot control samples in the PRJNA382490 (heavy metals) dataset we identified 95 genes in shoots that used 40% of the TPM values (396,812 TPM) compared to the same genes in roots (0.02%, 209 TPM). Two RBCS1A genes (EoRNA2_chr5H054244, EoRNA2_chr5H054248) and RuBisCO activase (EoRNA2_chr4H040697) alone use 8.5% of the allocated TPM values. These extreme differences mean that there are more TPM values to proportionately allocate in non-photosynthetic tissues/organs that do not express these high expressing genes. Genes that may be expected to be expressed consistently across a broad range of transcriptionally active cells include splicing factors such as U1A (EoRNA2_chr1H002474) and U1C (EoRNA2_chr6H067504), which showed 2-3 times the levels of TPM values in the root compared to the shoot. This difference may be the result of a combination of variable expression between the tissues and/or the increased TPM values associated with the non-photosynthetic root tissue (Fig. 3B).

**Figure 3.**
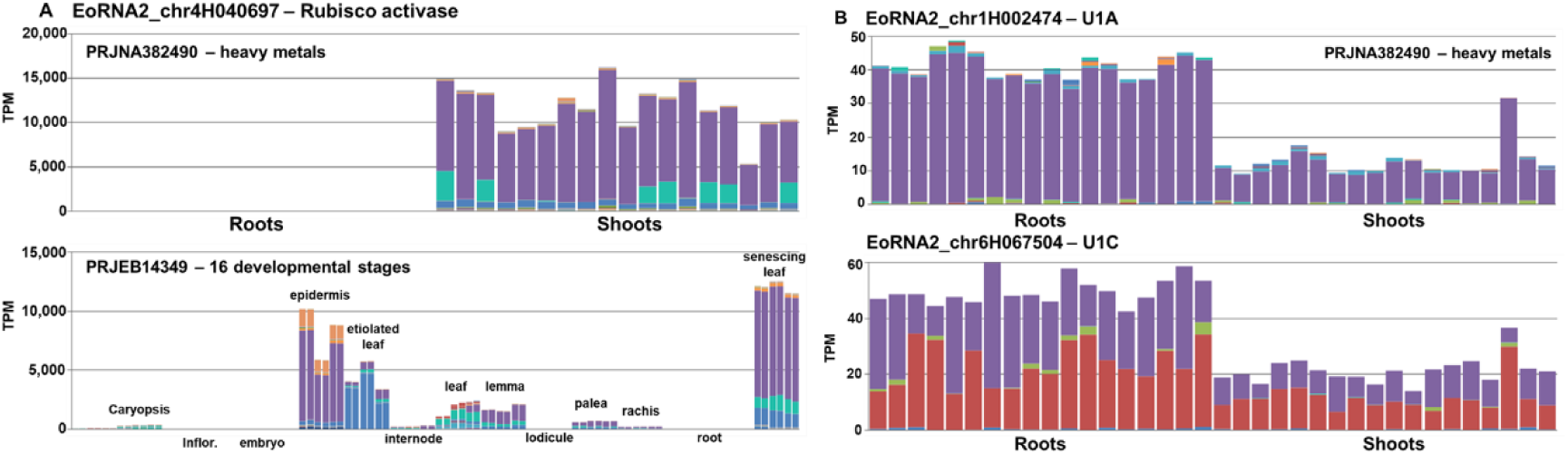
Expression differences between photosynthetic and non-photosynthetic tissues. A. EoRNA2 display of the photosynthetic gene RuBisCO activase (EoRNA2_chr4H040697) in two different studies, one comparing shoot and root (PRJNA382490) and another comparing a range of 16 different tissues and organs (PRJEB14349). B. Expression of two splicing factor genes U1A (EoRNA2_chr1H002474) and U1C (EoRNA2_chr6H067504) comparing shoot with root (PRJNA382490). Each bar represents a single biological repeat with the TPM values on the Y axis and the tissue displayed on the X-axis.

Ultimately, RNA-Seq TPM values represent relative transcript abundance within each sample rather than absolute expression levels. Gene-level TPM was obtained by summing the TPM values of all transcripts for each gene. TPM normalisation corrects biases caused by read depth and transcript length. However, further normalisation measures across samples/experiments will require assumptions (e.g. consistent expression across all datasets, see below).

We investigated different approaches to normalising the extreme differences in TPM between photosynthetic and non-photosynthetic samples. One possibility was to remove these 95 genes from the RTD and recalculate the TPM values. Although this could reduce the dominance of these transcripts, it would require that the set of dominant transcripts is known *a priori* for every tissue type and experimental condition, which is not feasible. This approach also invalidates the aim of the database to present the global expression levels of all the genes and transcripts within an RTD in the different samples. The second approach was to apply median normalisation to the dataset. The median is a robust measure for normalisation of the centre of the abundance distribution across transcripts and less affected by outliers. Median normalisation forces the median expression to be the same for all samples. As a test, we utilised the PRJNA382490 (heavy metals) dataset, because this contains three treatments over three replicated experiments in non-photosynthetic roots and photosynthetic shoots. Applying median normalisation across these samples minimised the sensitivity to the extreme values (Figure 4A). As expected, median normalisation did not alter the extreme differences found for the highly expressed photosynthetic genes (e.g. RuBisCO activase, EoRNA2_chr4H040697; Figure 4B). For some genes, median normalisation normalised the data across all the samples (e.g. RS31 splicing factor, EoRNA2_chr4H047631; Figure 4C), but in many cases median normalisation had a minimal effect and did not improve data analysis, particularly in shoot samples that were close to zero TPMs (e.g. Plant Cadmium resistance 2, EoRNA2_chr5H061177; Figure 4D). We therefore concluded that applying a median value to all the datasets to allow a more stable visualisation of the data did not enhance the data as presented by TPM.

**Figure 4.**
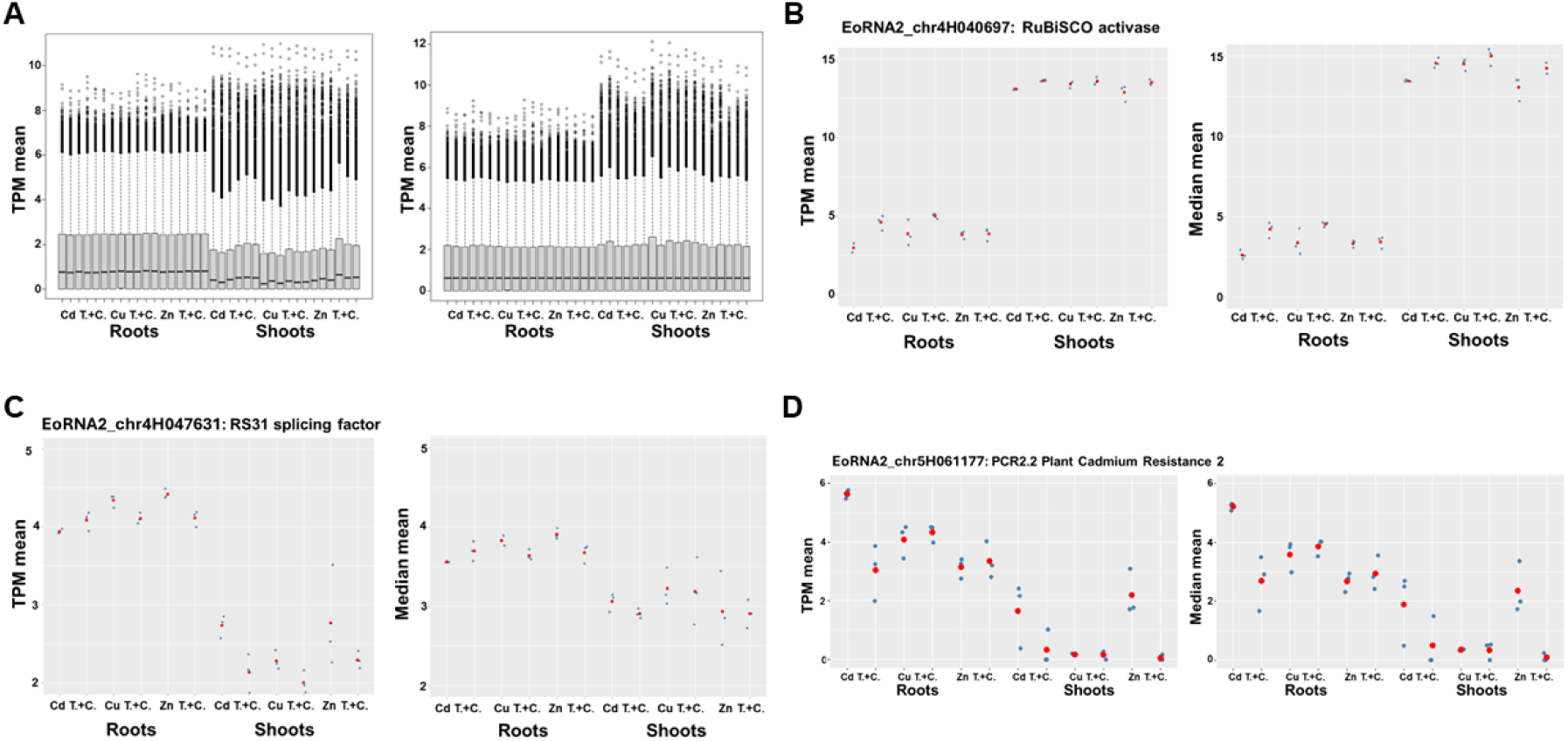
Application of median statistics to one EoRNA2 study dataset to test the normalisation of photosynthetic and non-photosynthetic tissues and organs. A. The logTPM values across the whole dataset compared to the logTPM after application of the median value. Subfigures B-D represent the means of the logTPM values of three different genes before and after application of the median value. B. The photosynthetic gene RuBisCO activase. C. The consistently expressed RS31 splicing factor. D. The Cd and Zn specific response gene PCR2.2. Cd represents the Cadmium treatment (T) and its control (C). Cu represents the Copper treatment (T) and its control (C), Zn represents the zinc treatment (T) and its control (C) along the X-axis.

Finally, we attempted to normalise the datasets using established housekeeping genes HvPP2AA2, Protein Phosphatase 2A subunit (EoRNA2_chr5H054343) and HvUBC21, ubiquitin-conjugating enzyme (EoRNA2_chr5H058172), which were used for quantitative PCR analyses^76^. The use of these genes assumed that their expression remains relatively constant across all the samples and datasets. To minimise potential complications arising from gene copy number variation, we selected genes that are present as single copies in the genome. Using PRJNA382490 (heavy metals) as an example dataset, we divided the housekeeping gene TPM value (*TPMh*) for each sample by the TPM value for the target gene (*TPMt*) for each sample (*TPMt*/*TPMh* = *TPMn*). In this selected dataset, the application of the selected housekeeping genes successfully improved the observed expression of the splicing factor U1A (and other splicing factors - not shown), which was expected to be consistently expressed in growing and dividing tissue/organs. This did not alter the significant differences in expression found in the photosynthetic genes or the differences in genes that respond to the heavy metal treatment application (Fig. 5), suggesting this as a suitable method to compare and quantify samples directly in this experimental dataset.

**Figure 5.**
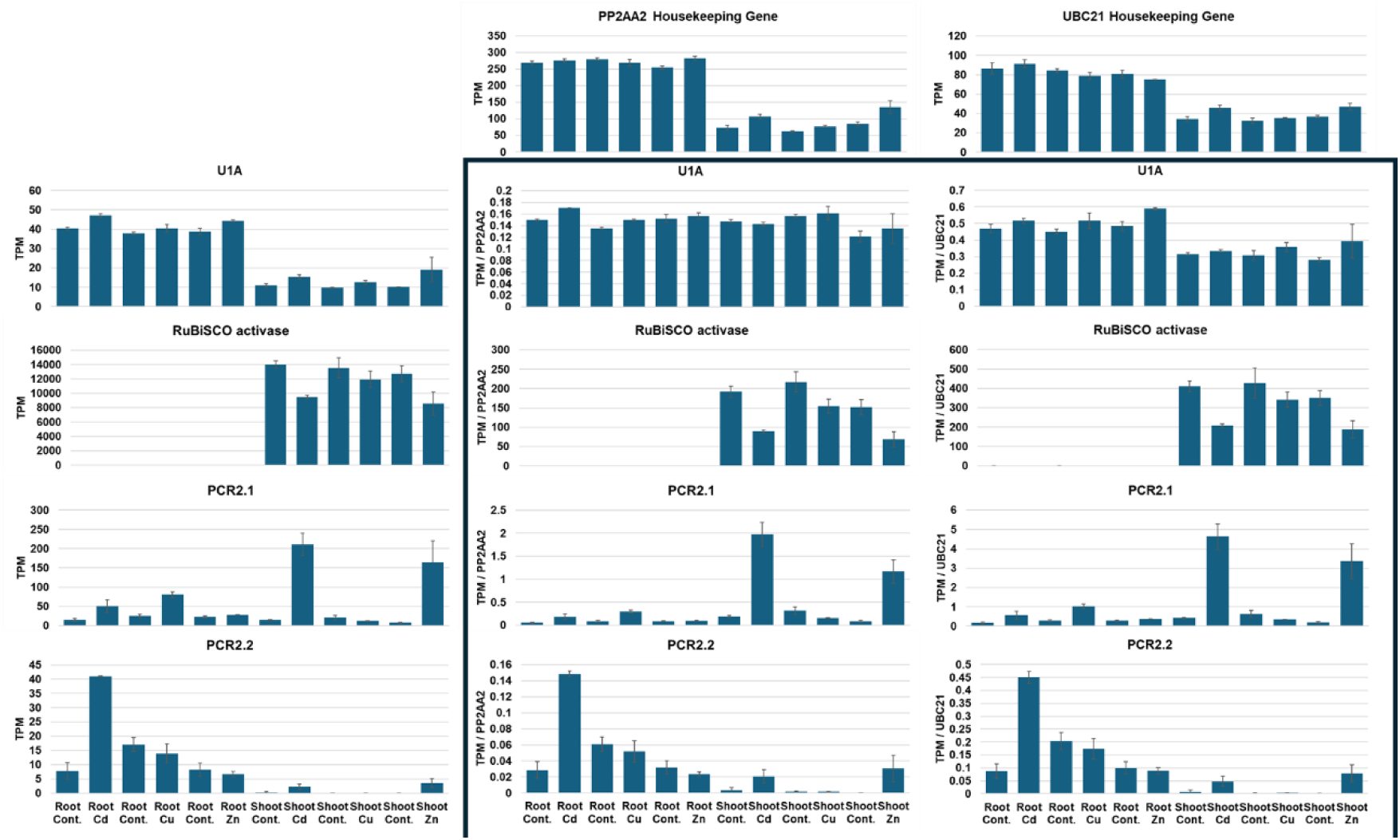
Normalisation of target gene expression by application of housekeeping genes. The two graphs along the top represent the expression in TPM for the housekeeping (HK) genes, PP2AA2 (EoRNA2_chr5H054343) and UBC21 (EoRNA2_chr5H058172). The graphs on the left show the expression of four target genes (U1A, RuBisCO activase, PCR2.1 and PCR2.2). The graphs in the box represent the normalisation of target gene expression by the HK genes, given as TPMt/TPMh. Each graph bar represents the mean of three biological reps with standard errors. The samples indicate root and shoot samples and the treatments (Cadmium- Cd, Copper - Cu and Zinc – Zn) and their controls (Cont.).

However, when we explored expression of the housekeeping genes across the complete EoRNA2 dataset, we found sample sets where there was very low expression of the housekeeper genes, which invalidates the assumption of consistent expression and application of these genes as housekeepers across the whole 171 EoRNA2 study datasets, and would limit their application in these samples. Other genes such as Elongation factor EF-2 (EoRNA2_chr5H061285), Malate dehydrogenase (EoRNA2_chr1H004773) and actin (multiple copy genes, e.g. EoRNA2_chr1H000339) have also been described as stably expressed genes across a range of tissues and could be applied to this type of normalisation^77^.

The EoRNA2 database and website, therefore, solely provides the relative TPM values for each dataset, that allows the researcher to perform their own quantifications and statistical analysis of the data. We do not correct for batch effects, outliers between samples or determine significant differences among experimental studies or changes because of scale, such as that observed between photosynthetic and non-photosynthetic tissues. The analyses described here are intended to make the researcher aware that these differences can exist between sample sets (we chose an extreme case to illustrate) and that analysis should be performed by the researcher depending on the sample studies selected. EoRNA2 facilitates these studies and we recommend systematic DEG and DAS expression analysis that utilises tools such as EdgeR^78^, Limma Voom^79^ or DESeq2^80^, wrapped up in user friendly commercial and/or academic, web based platforms such as Galaxy^81^, DEApp^82^, RaNA-seq^83^, iDEP^84^, basepair (basepairtech.com), 3D RNA-Seq^85^ (SHARP-GA), and THRAISE^86^ to describe differential gene expression and alternative splicing on the basis of fold differences with significance testing to identify the relevant genes responding to tissues or conditions across contrasting samples.

### Tissue and condition specific expression

EoRNA2 displays the expression data from 171 separate studies, and many datasets utilise similar tissues or conditions. A search across multiple separate experiments can identify shared tissue or condition specific expression. To validate this approach, we identified genes in EoRNA2 that have known tissue and condition specificity. *CYP704B* (EoRNA2_chr4H046163) has been described as having a vital role in anther development and is associated with male sterility^87^. EoRNA2_chr4H046163 shows exclusive high expression in anthers and with associated spike and inflorescence tissues (Fig. 6A). *GA2ox7* (EoRNA2_chr3H027484) is predominantly expressed in seed and required to deactivate gibberellin and maintain seed dormancy. In EoRNA2 this gene is almost exclusively expressed in developing seed and grain studies across all 171 studies (Fig. 6B).

**Figure 6.**
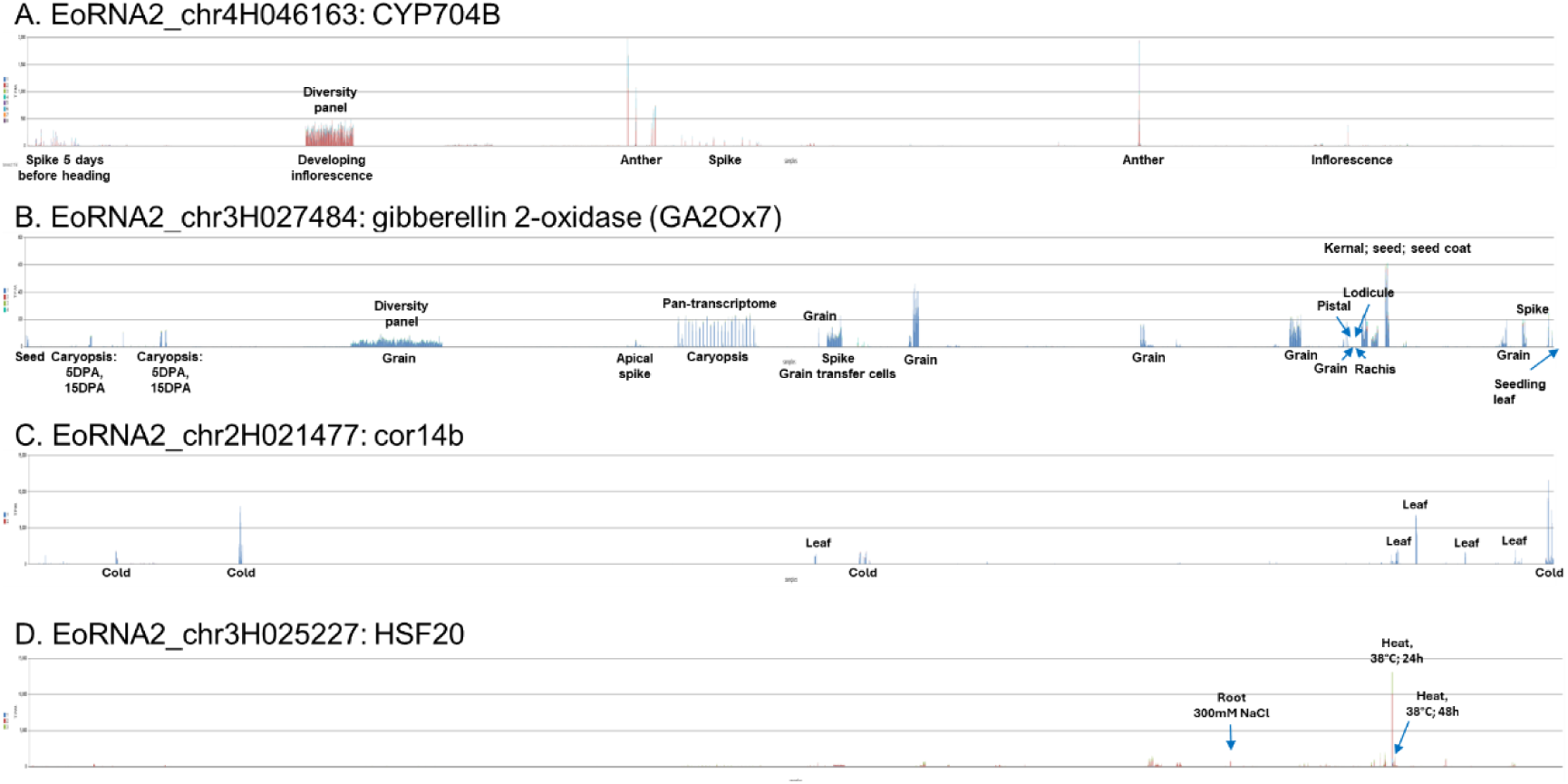
Tissue and condition specific gene expression. Graphical representation of the expression of 4 genes that show tissue and condition specific expression across all 171 EoRNA2 studies. The x-axis represents all 6,285 RNA-Seq samples in EoRNA2 and the y-axis represents the TPM value. Each peak represents one or more bars in the bar plot showing induction of expression in one or more samples. Selected experimental studies are labelled and arrowed. See the text for the description of the 4 genes.

Similarly, EoRNA2 shows conditional response genes. For example, Cor14b (EoRNA2_chr2H021477) is a low temperature and high-light specific responsive gene (Fig. 6C) and HSF20 (EoRNA2_chr3H025227) is highly and transiently expressed compared to all other tissues at 38°C for 24h (Fig. 6D). In this final example, there is expression in seed/grain and in response to other stresses such as 300mM NaCl in root tissue, but the difference represents a 17-fold variance in expression levels compared with high temperature. This confirmatory approach between different datasets represented in EoRNA2 provides confidence that a gene and its transcripts show true tissue/condition specific expression.

### Transcript variation

Most expression atlases focus on gene-level expression only. EoRNA2’s stacked expression graphs show the expression of the individual transcripts that make up the total gene expression value. For such a broad array of different genotypes, tissues, organs and conditions, presentation of the different transcript expression levels in EoRNA2 allows the researcher to identify the important gene transcripts in the expression analysis across all the study datasets. Transcript variants in EoRNA2 may be represented through alternative splicing, alternative transcript starts, alternative polyadenylation and through genotype variation. These transcript variants are further visualised in the gene models represented in the EoRNA2 JBrowse feature^59^, or by aligning the transcript sequences displayed in EoRNA2 to genomic sequences available through Ensembl (https://plants.ensembl.org/Hordeum_vulgare/Info/Index) and GrainGenes (https://graingenes.org/GG3/genome_browser).

### Genotype variants

In the first version of EoRNA we presented cultivar variation in the splicing factor gene RS31 (BaRT1_u-06868)^1^. The variation was represented by changes in the number of GCAG repeats at the end of an intron/start of an exon in the 5’ untranslated region (UTR) of the gene. EoRNA2 shows the same gene represented by EoRNA2_chr2H010834, (Fig. 7), and the new database contains 17 diversity studies, which allows further genotype transcript variants to be identified over a broader range of cultivars, landraces, wild and mutant genotypes. RS31 expressed by the genotype WI4330 shows 2 copies of GCAG compared to the previously described cultivars Clipper (4 copies of GCAG) and Sahara (3 copies of GCAG) (Fig. 7A, B). The many recently sequenced barley genomes may be used to further validate and confirm these genotype differences (e.g. barley pan-genome v.2 with 76 sequenced accessions^13^, https://graingenes.org/GG3/genome_browser). The diversity panel showing 209 genotypes registered for the UK market^88^ shows the diversity between the cultivars for transcripts 6 and 10 (Fig.7 C). The Morex genotype shows differential gene expression across 16 different tissues specific to transcript 6 (Fig.7 A), supporting a genotype specific transcript rather than, for example, tissue specific alternative splicing. Genotype transcript variants may also be the result of mutations at splice sites or sequences that support or inhibit splice site selection and result in genotype specific spliced transcripts that contribute to evolving genotypic transcript variants in EoRNA2^89^. Examples of genotype specific spliced transcripts have been previously shown in the barley pan-transcriptome^40^.

**Figure 7.**
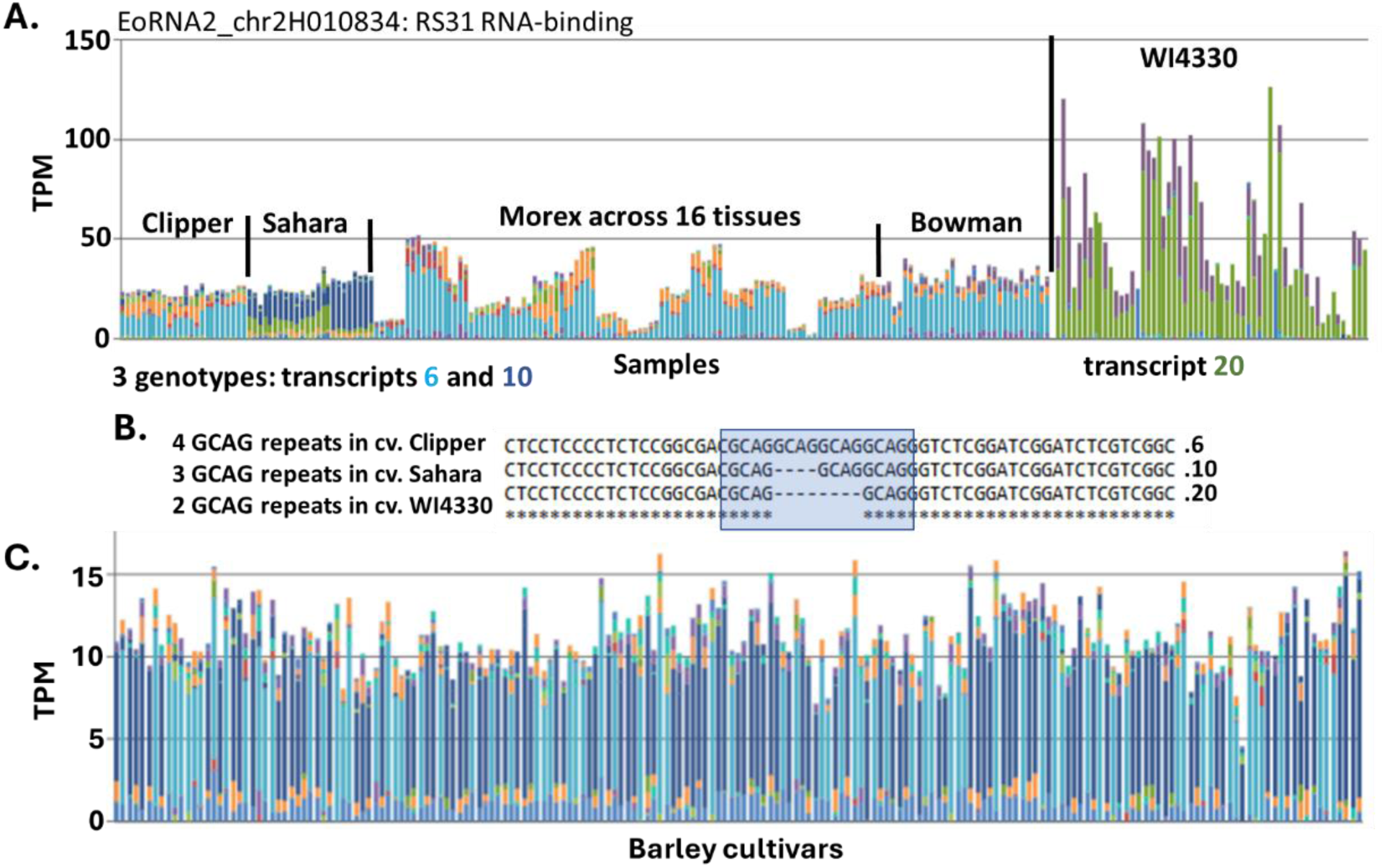
Genotype variant transcripts of the RS31 splicing factor gene. A. Genotype transcript variation between 5 genotypes showing, largely, transcripts, 6 (light blue), 10 (dark blue) and 20 (green). The Morex genotype shows differential gene expression across 16 different tissues to genotype specific transcript 6. B. Transcript sequence alignment showing the GCAG number variation between transcripts 6, 10 and 20. C. Transcript 6 and 10 variation between 209 barley cultivars in the crown tissue. In A and C the samples are arranged along the x-axis and expression levels given as Transcripts Per Million (TPM) along the Y-axis.

### Alternative Splicing

Differential alternative splicing is a vital part of regulatory gene expression^68^ ^90^. Some transcript expression changes in alternatively spliced transcript range from small but significant changes in transcript levels to cases where alternative transcripts completely switch abundances in response to a developmental stage or as a response to a stress condition^72^. EoRNA2 displays these changes across tissues, organs, stresses and genotypes. The splicing factor U2AF35A (EoRNA2_chr5H056610) is represented by 17 transcripts in EoRNA2, but only 3 of these transcripts (transcripts 8, 10 and 12) were transcribed and alternatively spliced to any great level of expression in 16 developmental stage tissues (Fig. 8A). Translation of these three transcripts identified transcripts 8 and 10 as having 5’UTRs that contain upstream open reading frames (uORFs), which may alter translation efficiency^91^. Transcript 8 also spliced a CDS internal intron within the coding sequence that leads to a shortened translatable protein at the C terminus of the protein. Only transcript 12 was predicted to generate a transcript with a protein coding sequence free of uORFs, through selection of an alternative downstream 3’ splice site that removed the uORF with the intron. The three alternative transcripts show variation in their TPM values between the different tissues, highlighting alternative transcript variation and its possible role in regulating the expression of genes in different tissues. For example, the tissues epidermis, etiolated leaf and senescing leaf were mostly represented by the uORF containing transcript 10 while developing inflorescence was mostly represented by both transcript 8 and 12.

**Figure 8.**
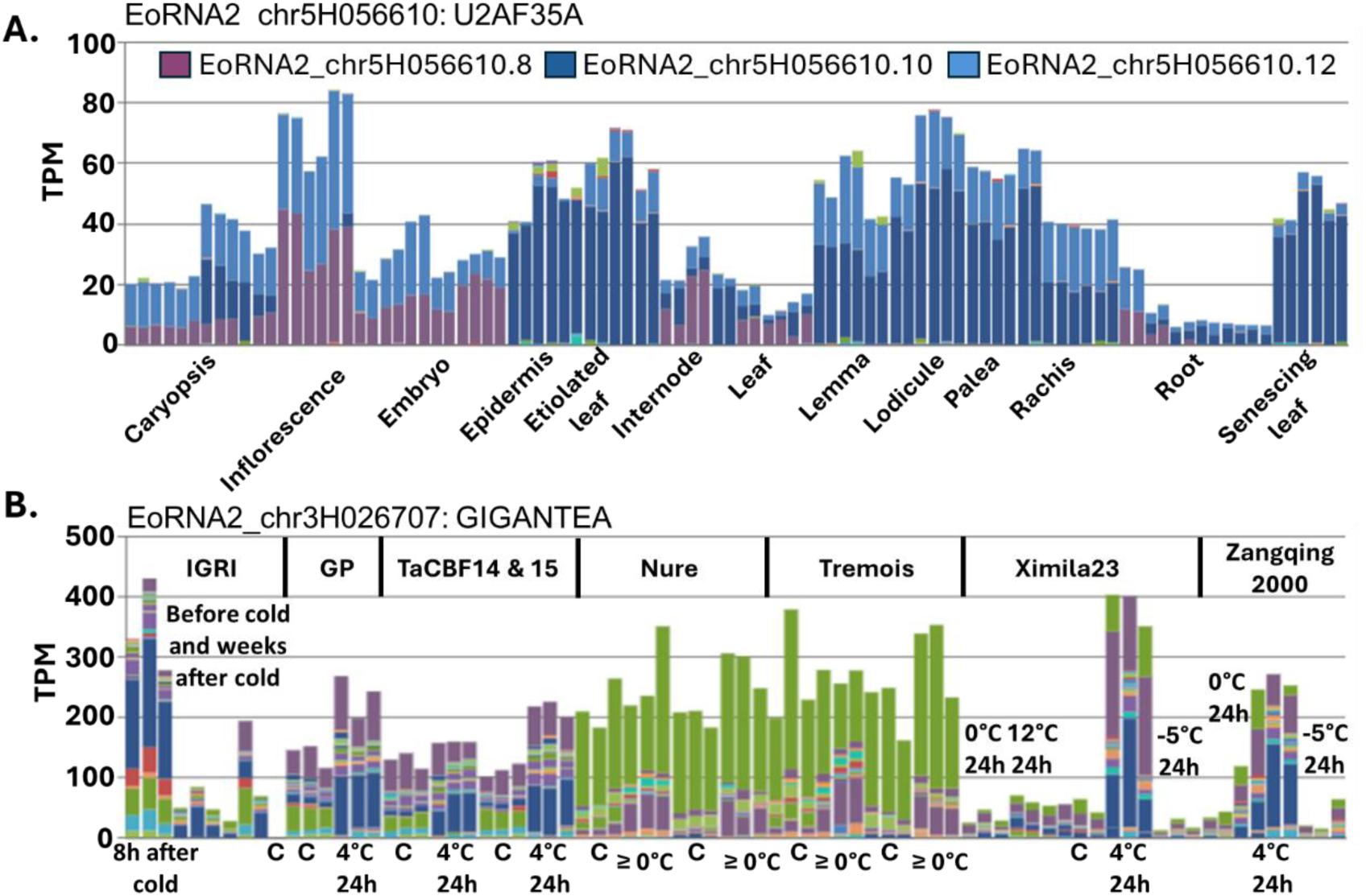
Alternatively spliced transcripts in barley tissues and low temperature conditions. A. Transcript expression of EoRNA2_chr5H056610/U2AF35A, across 16 developmental stages and tissues (PRJEB14349) showing three alternatively spliced transcripts. Samples are arranged along the x-axis. B. Transcript expression of EoRNA2_chr3H026707/GIGANTEA in 4 separate experimental studies of low temperature (PRJEB18276, PRJEB46519, PRJNA309882 and PRJNA999383). Control (C) and 4°C samples are labelled along the x-axis. Barley genotypes and transgenic barley types used are indicated at the top. GP is Golden Promise. TaCBF14 and 15 are two transgenic plants expressing wheat CBF genes. Expression levels in both graphs are given as Transcripts Per Million (TPM) along the Y-axis.

Similarly, alternative splice selection can occur as a response to different environmental conditions. GIGANTEA (GI, EoRNA2_chr3H026707), is a key circadian gene that plays a role in flowering time and the cold stress response in Arabidopsis. Gene expression is induced by cold stress, independent of the C-repeat Binding Factor (CBF) cold response pathway and the late flowering gi-3 mutant is sensitive to freezing^92,93^. A screen of all EoRNA2 studies identified an undescribed alternative splicing event that occurred under low temperature. Six studies in EoRNA2 involve reduced temperature and using the available metadata table further reduced this to four studies^94–97^ that were specific to replicated cold/freezing studies to investigate this alternative splicing event further. One of the studies also contained a subset of the study data relevant to drought, which were further excluded to leave only the low temperature data. Transcript analysis of these studies showed three main transcripts. Transcript 9 contained an exon within the 5’UTR that included an uORF. Transcript 74 was the result of an alternative transcript start (see below) that may lead to translation of a truncated version of GI. Transcript 21 was the result of an exon skip in the 5’UTR that removed the uORF sequence to produce a transcript with the protein coding sequence, unconstrained by an uORF. This alternatively spliced transcript is specific to a short-term response to low temperature (4°C), above freezing temperatures and below higher temperatures (12°C) as shown by the different experimental studies in EoRNA2. In this case, low temperature at 4°C increased gene transcription and produced a transcript that lacks any inhibition to translation. EoRNA2 therefore allows the researcher to filter and select their specific tissue/organ/condition relevant to their study. It further generates new gene regulation hypotheses, which can be further validated; in this example the significance of the alternative splicing regulation found in *GI* may be tested further in relation to low temperature and circadian changes by molecular techniques.

### Transcription start sites (TSS) and alternative polyadenylation(APA)/transcription end sites (TES)

5’ and 3’ UTRs affect transcript localisation and are sites for regulatory RNA binding proteins. They also affect mRNA stability through nonsense mediated decay (NMD) or miRNA targeting sites, and translation efficiency through uORFs, which also have a role in NMD^91,98–102^. The length of 5’ and 3’UTRs are variable because of alternative splicing, alternative polyadenylation sites in the 3’ UTR and through alternative transcription start and termination sites. The BaRTv2 reference transcript dataset contains transcripts with supported transcript start and end sites and is incorporated into the EoRNA2_RTD. PanBaRT20_chr5HG50838 (formate tetrahydrofolate ligase ortholog; EoRNA2_chr5H055935) was recently described to have transcripts with an alternative TSS that extended the 5’UTR specifically in caryopsis^40^. Other transcripts result in a shortened CDS and 5’UTRs with an uORF. That study (PRJEB64639) and these transcripts are found in EoRNA2 and can be further studied across the range of all the samples and conditions presented in EoRNA2. Examples include transcript 10 that splices a rare U12-dependent intron (AT_AC) and transcript 16 with a TSS that leads to a shorter transcript and a shorter CDS. Similarly, alternative polyadenylation leads to the formation of transcripts with alternative 3’UTR lengths. EoRNA2_chr2H010731 represents a transcription initiation factor that shows alternative TESs between transcripts 2 and 4 in different tissues without alteration of the CDS (see PRJDB14465 as an example). The effect of these alternative 3’ ends remain to be determined in this gene and studied further.

### Working Example - Cleistogamy

New areas of investigation may be developed through an understanding of the gene and transcript expression profiles displayed in EoRNA2. For example, important genes may be identified through insights provided by studying the expression of the genes found between two molecular markers that are associated with a particular trait. Differential gene and alternative splicing analysis can reveal genes that are expressed in a tissue- or condition-specific manner associated with the marker and phenotype. Barley mutant populations can also help identify genes that are relevant to a tissue specific trait or conditional response to a stress, and some of these mutant populations have been sequenced and are found within EoRNA2. Many orthologous genes are characterised in related species through mutational analysis, and the expression of these genes can help support their translation into barley. For example, cleistogamy allows self-fertilisation of flowers, promotes seed set and prevents dissemination of pollen in barley due to a closed flower phenotype. This phenotype in barley is the result of poorly developed lodicules, the result of a mutation in the miRNA target site in the 3’ UTR of the barley *Cly1* gene (Genbank: KF261344)^103^. *Cly1* gene expression in the EoRNA2 database is complex, it has many transcripts and is expressed across a broad range of tissues and organs as well as the lodicule (EoRNA2_chr2H022984) affected by the trans-acting miRNA. The rice cleistogamous mutant *superwoman1-cleistogamy1* (*spw1-cls1*) has an I45T mutation in OsMADS16 that leads to reduced interaction with two other MADS box genes, OsMADS2 and OsMADS4^104,105^. Orthologues of all three MADS box genes were identified in barley (OsMADS16 ≡ EoRNA2_chr7H083714; OsMADS2 ≡ EoRNA2_chr3H034592; OsMADS4 ≡ EoRNA2_chr1H007161). Expression analysis displayed in all three genes was highly specific to the lodicule (Fig. 9) with some reduced expression displayed in anther, stigma, developing inflorescence, early caryopsis and spike tissues. These three barley gene orthologues are candidates for analysis of lodicule development and cleistogamy by finding available barley mutants or using barley gene editing^106,107^. EoRNA2 provides transcript sequence information that allows the design of primers and/or editing targets to select or produce barley mutants in this key barley phenotype.

**Figure 9.**
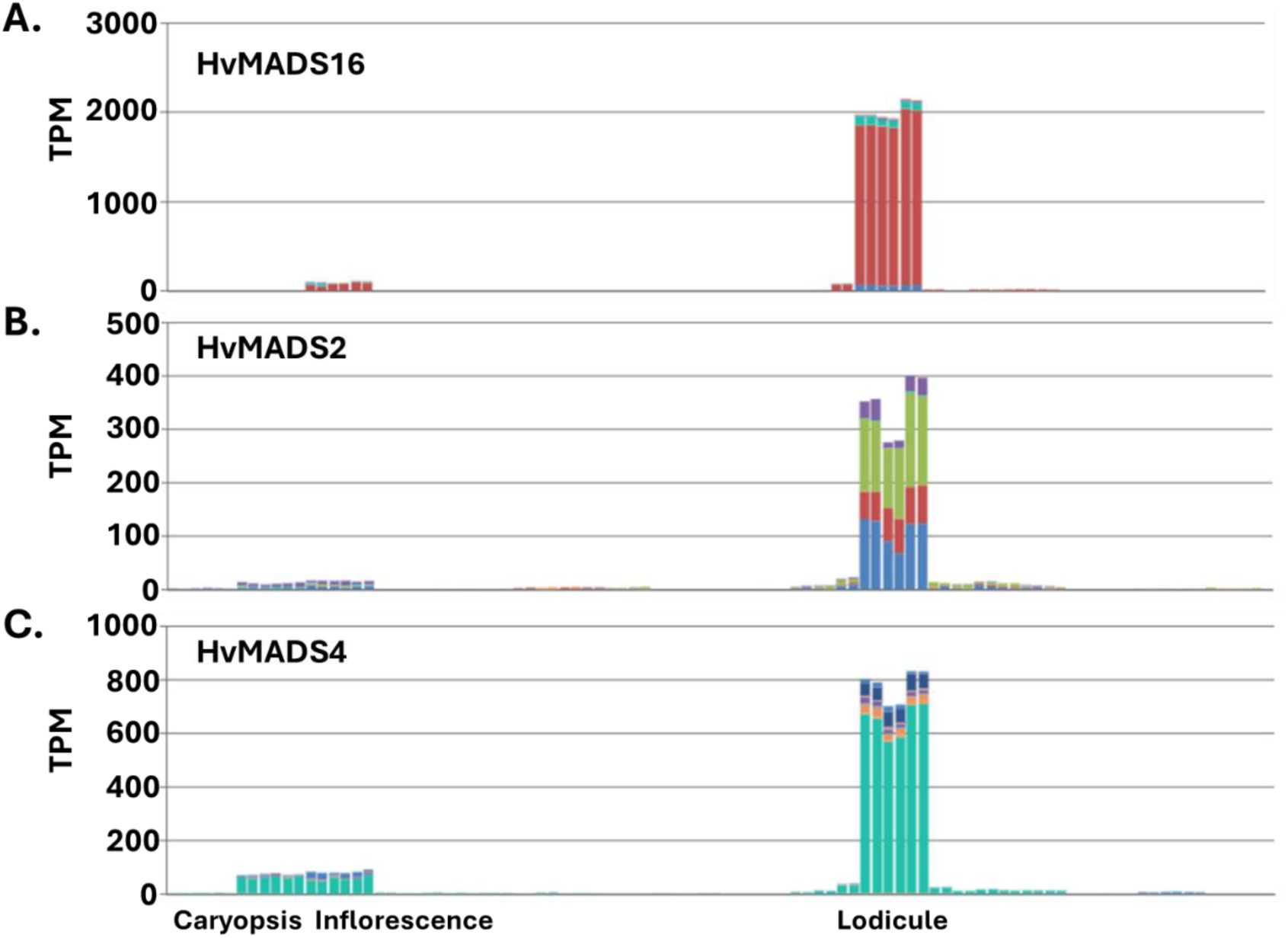
Lodicule specific expression of three barley MADS box genes. A. Expression pattern for HvMADS16. B. Expression pattern for HvMADS2 C. Expression pattern for HvMADS4. The samples from 16 developmental stages (PRJEB14349) are arranged along the x-axis and expression levels given as Transcripts Per Million (TPM) along the Y-axis. Go to EoRNA2 to download expression across all 171 barley studies (https://ics.hutton.ac.uk/eorna2/index.html).

### Summary and Future

The data displayed in EoRNA2 provides insight into barley biology through gene and transcript expression across a broad range of tissues. It is a constructive tool that helps generate hypotheses and questions specific to a researcher’s gene or genes of interest. EoRNA2 further links the expression information to genomic information. EoRNA2 is format ready for future single cell transcriptomes, poly-ribosome isolated transcriptomes and proteomic data. These extensive data sets can help train artificial intelligence (AI) systems that model the complexities of multi-transcriptomic data sets across thousands of genes and support AI guided gene editing prediction tools such as CRISPR_GPT^108^.

## Data Availability

The EORNA v.2 web interface is available at https://ics.hutton.ac.uk/eorna2/index.html. Tables with gene and transcript level TPM data and metadata, as well as the EORNA2_RTD files (transcript and protein FASTA files, GTF file, annotation, pan-genome reference) have been deposited in the Zenodo data repository under DOI https://doi.org/10.5281/zenodo.18466205.

## Code Availability

All code described in the manuscripts is available freely from https://github.com/cropgeeks/eorna-v2. This includes scripts for the creation of EORNA2_RTD, the Nextflow workflow for data discovery and quantification, as well as all scripts for setting up the database, importing data and creating the web interface.

## Author Contributions

LM developed the EORNA2 database, web interface and data import workflow and populated the database with the new TPM dataset. IM constructed the Docker image for deployment of the database and web server. CS conducted the data validation and developed the biological use cases. LM and CS jointly curated the metadata. WG developed the EORNA2_RTD. CDM conducted the statistical analyses involving data normalisation. MB developed the data discovery and processing workflow and applied it to the public barley data. CS and MB jointly developed and supervised the project. All authors contributed to writing and editing of the manuscript.

## Acknowledgements

We wish to acknowledge all the researchers who deposited their barley short RNA-Sequence data in the public databases. In the text we have utilised specific datasets indicated by their study accession number to illustrate different aspects of the database and website. We utilised these datasets because of the coverage of tissues and genotypes and because the experimental design of the study allowed us to illustrate our points easily and simply.

